# RBC-GEM: a Knowledge Base for Systems Biology of Human Red Blood Cell Metabolism

**DOI:** 10.1101/2024.04.26.591249

**Authors:** Zachary B. Haiman, Angelo D’Alessandro, Bernhard O. Palsson

**Affiliations:** Department of Bioengineering, University of California San Diego, La Jolla, California, United States of America; Department of Biochemistry and Molecular Genetics, University of Colorado Anschutz Medical Campus, Aurora, CO, USA; Department of Pediatrics, University of California San Diego, La Jolla, California, United States of America; Bioinformatics and Systems Biology Program, University of California, La Jolla, San Diego, CA, United States of America

**Author notes:** Corresponding author (BOP).

## Abstract

Advancements with cost-effective, high-throughput omics technologies have had a transformative effect on both fundamental and translational research in the medical sciences. These advancements have facilitated a departure from the traditional view of human red blood cells (RBCs) as mere carriers of hemoglobin, devoid of significant biological complexity. Over the past decade, proteomic analyses have identified a growing number of different proteins present within RBCs, enabling systems biology analysis of their physiological functions. Here, we introduce RBC-GEM, the most extensive and meticulously curated metabolic reconstruction of a specific human cell type to-date. It was developed through meta-analysis of proteomic data from 28 studies published over the past two decades resulting in a RBC proteome composed of more than 4,600 distinct proteins. Through workflow-guided manual curation, we have compiled the metabolic reactions carried out by this proteome. RBC-GEM is hosted on a version-controlled GitHub repository, ensuring adherence to the standardized protocols for metabolic reconstruction quality control and data stewardship principles. This reconstruction of the RBC metabolic network is a knowledge base consisting of 718 genes encoding proteins acting on 1,590 unique metabolites through 2,554 biochemical reactions: a 700% size expansion over its predecessor. This reconstruction as an up-to-date curated knowledge base can be used for contextualization of data and for the construction of a computational whole-cell model of a human RBC.

**Author Summary:** Human red blood cells (RBCs) have been studied for decades because of their unique physiology, essential oxygen delivery functions, and general accessibility. RBCs are the simplest yet most numerous of human cell types due to the loss of cellular organelles during their development process. This process has evolved to maximize hemoglobin content per cell to facilitate RBCs’ main function in gas transport. RBCs are integral to a variety of medical applications, such as blood storage for transfusion. Recent advancements in high-throughput data collection have greatly expanded our understanding of RBC metabolism, highlighting important roles and functions for RBCs in maintaining homeostasis in the organism in addition to oxygen transport. Here we provide a knowledge base for the human RBC as a genome-scale metabolic reconstruction. Our results highlight the complexity of RBC metabolism, supported by recent advancements in high-throughput data collection methods for detecting low-abundance proteins in RBCs. We make knowledge about the RBC findable, accessible, interoperable, and reusable (FAIR). As RBC research is likely to see many translational medical advancements, a knowledge base for the contextualization of RBC data will serve as an essential resource for further research and medical application development.

## Introduction

Recent estimates suggest that red blood cells (RBCs) are by far the most numerous cell-type in the human body, accounting for ∼83% of total human cells in an adult. The average lifespan of a healthy human RBC is approximately 120 days, during which the RBC undergoes approximately 200,000 circulatory cycles[1]. Each transit cycle takes approximately one minute. RBCs thus are subjected to constantly changing environmental stresses that exacerbate damage and degradation of proteins, from elevated oxidant stress in the lung, to hypoxia and shear stress when traversing capillaries as narrow as 5 μm[2]. The RBC lacks the machinery necessary to synthesize new proteins *de novo*; consequently, the RBC proteome loses functionality over time, affecting essential functions in gas transport. Irreversible modifications to metabolic enzymes at key functional residues have been documented as a function of RBC aging in vivo and in vitro, ultimately promoting proteasome-dependent degradation of rate-limiting enzymes in key energy and redox metabolic pathways (e.g., glyceraldehyde 3-phosphate dehydrogenase – *GAPDH* or glucose 6-phosphate dehydrogenase – *G6PD*)[3–5]. Furthermore, both genetic and environmental factors influence irreversible glycation of hemoglobin, thus the level of glycated hemoglobin has become a clinically important biomarker for glycemic control[6–8].

The relative simplicity of the RBC with respect to other cell types, the lack of organelles, and its central, yet specialized role in systems physiology led to the consensus view that RBCs are inert cells with limited metabolic capabilities. RBC metabolism is tailored to sustain survival in circulation, deformability through the circulatory system, and, above all, the metabolic-dependent regulation of oxygen kinetics. As such, the focus on RBC metabolism has historically been limited to a subset of metabolic pathways, including energy metabolism via glycolysis, oxidative stress handling by the glutathione systems and the pentose phosphate pathway, and the allosteric regulation of hemoglobin oxygen-binding affinity through 2,3-bisphosphoglycerate (2,3-BPG). However, two decades of proteomic studies have now elucidated an unexpected complexity of the RBC proteome, prompting a reevaluation of the purported simplicity of RBC metabolism. Despite a wealth of accumulating data, a systematic review of the literature and its organization according to latest standards in the field of systems biology is currently missing, creating the impetus for the current study.

Genome-scale models (GEMs) of metabolism serve as a platform to integrate multiple biological data types that can subsequently be interrogated and interpreted within a metabolic context. Human GEMs contain all metabolic reactions known to occur across multiple human cell types, encompassing all possible metabolic genes defined in the human genome without regard for tissue or cell specificity[9,10]. Several cell-specific and/or context-specific human GEMs can be derived by mapping transcriptomic data or proteomic data onto these network reconstructions. GEMs derived from these data types are well-suited for the analysis of biological functions[11].

The first proteomically-informed metabolic reconstruction of the human RBC, iAB-RBC-283[12], was derived from the first global human reconstruction, Recon1[13]. The iAB-RBC-283 knowledge base has been successfully utilized in numerous applications, including the development of personalized whole-cell kinetic models of RBCs[14], the elucidation of temperature dependence of RBC metabolism[15], and the exploration of host-parasite metabolic interactions in *Plasmodium falciparum*-infected RBCs[16,17]. Several proteomics studies were utilized in the construction of iAB-RBC-283, with the largest study listing approximately 1,500 proteins identified in the erythrocyte[18–21]. Despite this expanded coverage, connectivity analysis of iAB-RBC-283 highlighted the need for targeted studies on the functionality of non-canonical citric acid cycles in the RBC, believed to be inactive in enucleated RBCs[22]. Subsequent follow-up studies did indeed confirm the activity of several enzymes involved in citrate metabolism[23,24].

With the increasing sensitivity and accuracy of proteomic approaches, the number of proteins identified in RBCs has now grown to over 3,000[25,26], prompting the need to generate a new reconstruction that encompasses such substantial progress. Much of the metabolic complexity associated with RBCs is within the low-abundance proteome. Advances in protein quantification strategies, driven by the increased sensitivity of latest generation mass spectrometers and the development of novel bioinformatics tools, have made it possible to estimate the copy numbers of the low-abundance proteins[27,28]. The innovation and advances in affordable, high-throughput omics technologies continue to drive change in blood science and personalized transfusion medicine. These advancements promote the paradigm shift away from the long held views that RBCs are relatively inert cells[29–32], and “not a hapless sack of hemoglobin”, as Greenwalt put it[33]. Thus, there is a pressing need for an updated RBC knowledge base that reflects the last decade of advances made in RBC omics research. Furthermore, it is essential for improvements to be made in a tractable and transparent manner as new technologies and methodologies expand our knowledge of the true scope of RBC metabolism with immediate, translational implications for human biology.

Here we present RBC-GEM, an updated knowledge base of erythrocyte metabolism. The RBC-GEM knowledge base was developed by leveraging proteomic data from 28 publications and through manually curating decades of experimental literature primarily pertaining to human RBCs. We integrated the knowledge base with GitHub version control software and the MEMOTE suite for standardized GEM testing[34], adhering to both findability, accessibility, interoperability, and reusability (FAIR) principles for scientific data management[34–36] and minimum information requested in the annotation of biochemical models (MIRIAM) guidelines for annotation and curation of quantitative models[37]. RBC-GEM is one of the most comprehensive, curated reconstructions of a specific human cell type to-date. We describe key considerations throughout the iterative reconstruction process, explore the topology of the resulting metabolic network, and validate both presence and activity of enzymes through the multitude of collected evidence. Our goal was to construct a curated knowledge base for human RBC metabolism that could serve as a foundational framework for RBC research across a multitude of disciplines.

## Results and Discussion

### Generation of the RBC-GEM knowledge base

We began by reconciling the previously published iAB-RBC-283 reconstruction[12] with the current iteration of the Human-GEM reconstruction (version 1.18.0[38]). We then followed the protocols developed for generating high-quality GEMs[39], while adhering to guidelines set for previously developed version-control frameworks for rapid, trackable model updates of consensus GEMs[34–36]. The final reconstruction, designated RBC-GEM, was generated after subjecting the initial draft reconstruction to several refinement cycles guided by biochemical databases (Table A in S1 File), an abundance of publicly available proteomic data (Table B in S1 File), rigorous manual curation (Table C in S1 File), and repetitive assessments of model quality (Fig. 1).

**Fig 1.**
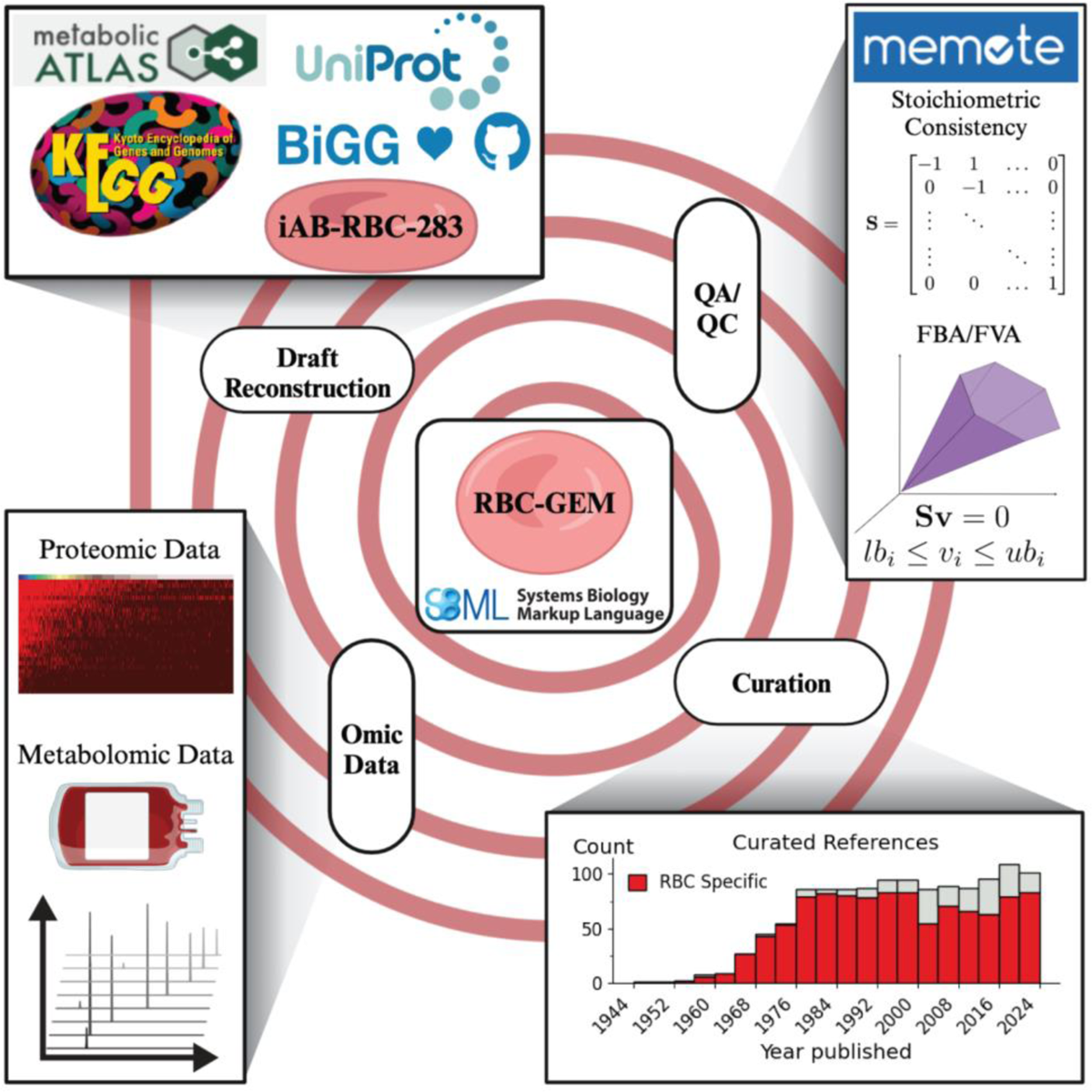
Workflow for the generation of the RBC-GEM knowledge base. The RBC-GEM was generated through several cycles of iterative expansion and refinement. For the first cycle, iAB-RBC-283[12] was downloaded from the BiGG Model database[40] as the initial reference network. New candidate reactions were identified by tailoring the global human reconstruction[38] for the RBC based on publicly available omic data, and by cross-referencing the data with biochemical reaction and protein databases (e.g., KEGG[41], UniProtKB[42]). The existence and catalytic activity of enzymes were confirmed through systematic manual curation [39] of experimental literature with the vast majority directly pertaining to human RBCs. The resulting knowledge base is set up within a version-controlled and open-source framework for tractable and traceable improvements, and was evaluated using the MEMOTE quality control (QC) utility [34]. Key database and software resources, collected proteomic data, and curated references are provided in the SI.

This study releases the first version of RBC-GEM (version 1.1.0). It represents the most comprehensive reconstruction of the RBC metabolic network to-date, representing a 700% size increase in reactions and 250% increase in genes compared to its predecessor (Fig. 2). This network contains 718 genes (Table D in S1 File), 1,590 unique metabolites (Table E in S1 File), and 2,554 biochemical reactions (Table F in S1 File). RBC-GEM represents the significant progress achieved towards the comprehensive elucidation of erythrocyte metabolism through integrated omics approaches, incorporating a total of 28 proteomic datasets published across the last 20 years (Table B in S1 File). RBC proteomic data is limited by challenges with sample purity as well as questions concerning relevance and activity of identified proteins[43]. We therefore used metabolomics data of RBCs across a variety of conditions to confirm the presence of metabolites and active pathways within the RBC[23,26,32,44–49]. Furthermore, we validated the reconstruction through extensive manual curation, culminating in a collection of over 1,000 publications specifically pertaining to human RBC metabolism (Table C in S1 File). RBC-GEM is the most comprehensive reconstruction of metabolism in the human erythrocyte, and the most detailed metabolic network in a human cell. The material in the SI archives the details of the reconstructed network.

**Fig 2.**
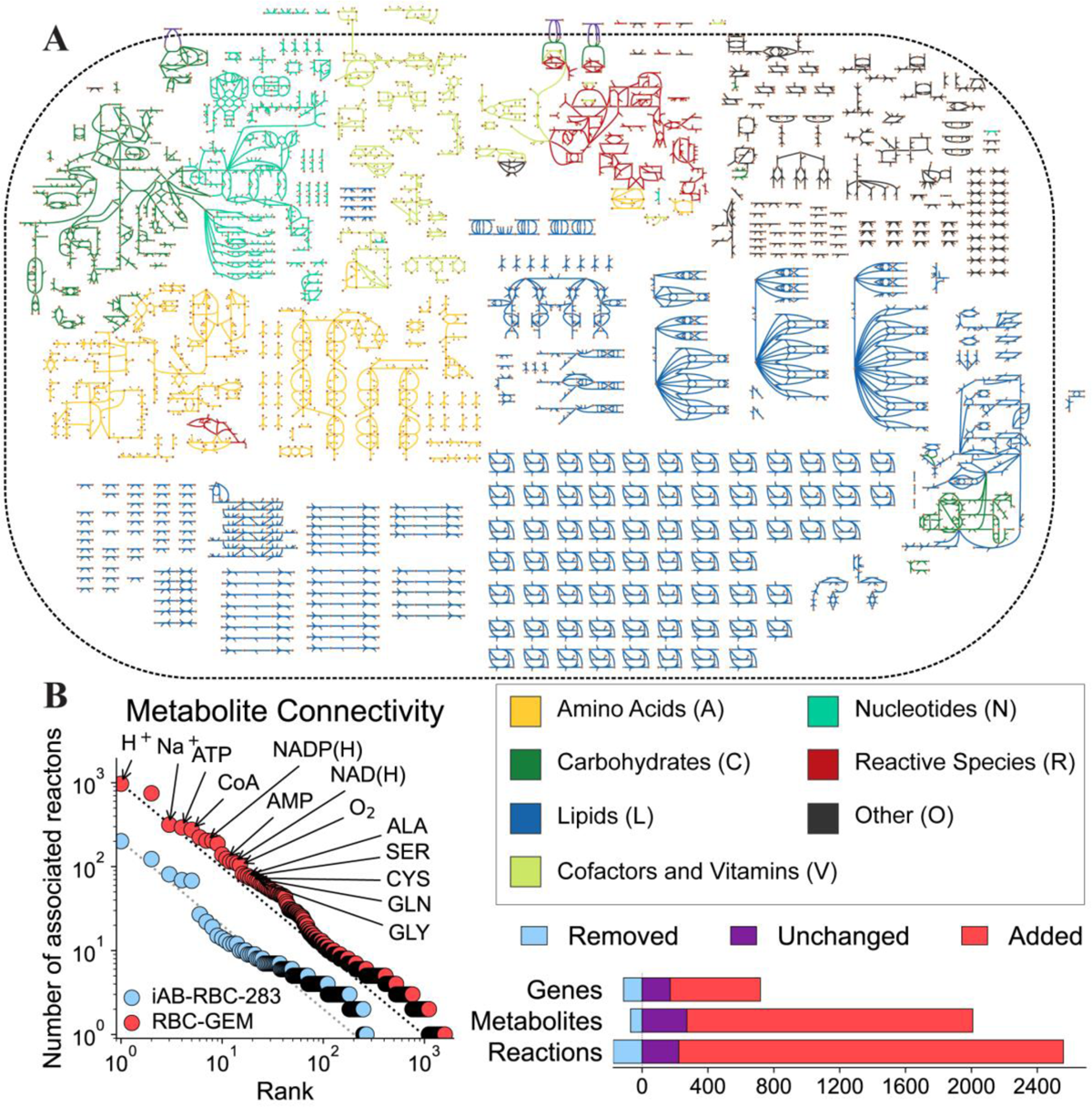
The expanded human RBC metabolic network. (A) The reconstructed RBC-GEM 1.1.0 metabolic network is larger in scope and complexity than all previous erythrocyte models. (B) RBC-GEM was derived through reassessment and systemic refinement of the iAB-RBC-283 [12] reconstruction components followed by the addition of several new components to expand the metabolic network based on new multi-omic evidence and backed by decades of RBC literature. The resulting RBC metabolic network has a high connectivity with many points above the reference line, reflecting a more interconnected RBC metabolic network with over twice the proteomic coverage compared to its predecessor. Transport reactions were not included for visual clarity. The network map was created using the Escher Network visualization tool[50].

### RBC metabolism is contextualized and explored through network visualization

In tandem with network reconstruction, we developed a network map of the full RBC metabolic network using the Escher Network visualization tool[50]. Visualization of biochemical pathways through detailed maps has enabled researchers to understand the conversion of metabolic species and coordination of enzymes within the cellular environment[51]. Through the Escher visualization tool, the entire RBC metabolic network can be visualized at once. We utilized KEGG pathways[41] to group the metabolic reactions into seven general categories (Table G in S1 File), which we subsequently visualized onto the network map (Fig. 2A). We provide an interactive version of the map in the SI, in which user-defined data can be overlaid onto the map, providing visual context for the exploration of the RBC-GEM reconstruction and interpretation of data. Additionally, the map is provided as an Escher JSON file and in standard SBGN and SBML layouts (S2 File), generated by EscherConverter tool (version 1.2). The accessibility of the map allows for the derivation of new user-generated pathway visualizations via Escher without having to start from a blank canvas. Furthermore, new maps can be deposited within the version control framework of RBC-GEM where they can be maintained and reused across studies, aiding in the process of consensus building with respect to RBC metabolism.

### Metabolic pathways are connected through key cofactor pools

We computed the connectivity for each metabolite with and without regard for compartment boundaries (Table H in S1 File). The most connected metabolites were the cofactors that act as currency metabolites for energy metabolism (ATP, ADP, AMP) and redox metabolism (NADP(H), NAD(H)). RBCs rely on the glycolytic, pentose phosphate, and purine salvage pathways to maintain critical cofactor pools. Additional computational and metabolomic studies demonstrated that the citric acid cycle enzymes such as isocitrate dehydrogenase (*IDH1*), malic enzyme (*ME1*), and malate dehydrogenase (*MDH1*) contributed to cofactor pools through regeneration of NAD(P)H and provision of intermediates to pyruvate kinase (*PKLR*)[23,24,31]. Conversely, numerous cellular processes depend on ATP hydrolysis for energy, while NAD(P)H is required for reduction of methemoglobin, the oxidized form of hemoglobin, through NADH-dependent cytochrome b5 reductase (*CYB5R3*) and NADPH-dependent flavin reductase (*BLVRB*), also known as biliverdin reductase B. As insufficient levels of ATP and NAD(P)H increase hemolytic propensity, understanding their connections to other metabolic pathways and the causes for their depletion is paramount for a multitude of biomedical research applications[31,52–57].

The expansion of lipid metabolism can be partly attributed to numerous, sparsely connected lipid species undergoing a similar set of reactions, transferring lipids between the highly connected Coenzyme A (CoA) and L-carnitine cofactors. The free and esterified lipid species form distinct, yet equilibrating pools that fuel and buffer phospholipid acyl-chain turnover as part of the Lands cycle[58–64]. The sodium ion (Na^+^) and amino acids have similar metabolite connectivities in both the intracellular and extracellular compartments, highlighting their involvement in membrane transport processes. Calculation of the gene connectivities (Table I in S1 File) confirmed the roles of the LAT1 heterodimer (SLC7A5 and *SLC3A2*), a remnant of erythropoiesis with SNPs significantly associated with RBC kynurenine levels [65], and y+LAT2 heterodimer (*SLC7A6* and *SLC3A2*), initially identified via RBCs of individuals with Lysinuric Protein Intolerance [66,67], in driving the exchange of amino acids across the membrane. As seen in other mammalian cell types, these transport proteins facilitate uptake of cationic and neutral amino acids into RBCs, which can be subsequently effluxed as a driving force for the Na^+^-dependent and Na^+^-independent uptake of other neutral amino acids [68].

### Increased connectivity

An open question from the reconstruction of iAB-RBC-283 was whether the low metabolite connectivity was a consequence of an inherently ‘fragmented’ erythrocyte network or incomplete proteomic coverage[12]. In RBC-GEM we added hundreds of new genes, metabolites, and reactions to the iAB-RBC-283 reconstruction, supported by new multi-omic and experimental evidence. We compared the metabolite connectivity of RBC-GEM to its predecessor by connecting the minimum and maximum connectivity in each distribution with a reference line (Fig. 2B). The distribution for iAB-RBC-283 was observed to reflect a less connected network, indicated by the number of points below the reference line[12]. Conversely, the distribution of connectivities for RBC-GEM reflected a network with higher connectivity as indicated by the number of points above the reference line. Furthermore, the distribution for RBC-GEM was shown to be greater than the distribution of iAB-RBC-283. The increased coverage of the RBC proteome in RBC-GEM is attributed to the utilization of multiple published proteomic datasets, many of which were published after iAB-RBC-283.

### Twenty years of proteomic data generation help define the reconstruction

Transcriptomic and proteomic data is often utilized for the development of context-specific metabolic reconstructions for various cell types[11,69]. However, as human RBCs are devoid of nuclei, proteomic evidence was essential for developing the metabolic reconstruction, providing direct evidence for the existence of proteins. We collected published proteomic data for erythrocytes from 28 datasets[18–20,26–28,70–90] that span 20 years of RBC proteomic research (Table C in S1 File). This wealth of proteomic data was mapped onto RBC-GEM to gauge the level of supporting proteomic evidence for each protein (Fig. 3A).

**Fig 3.**
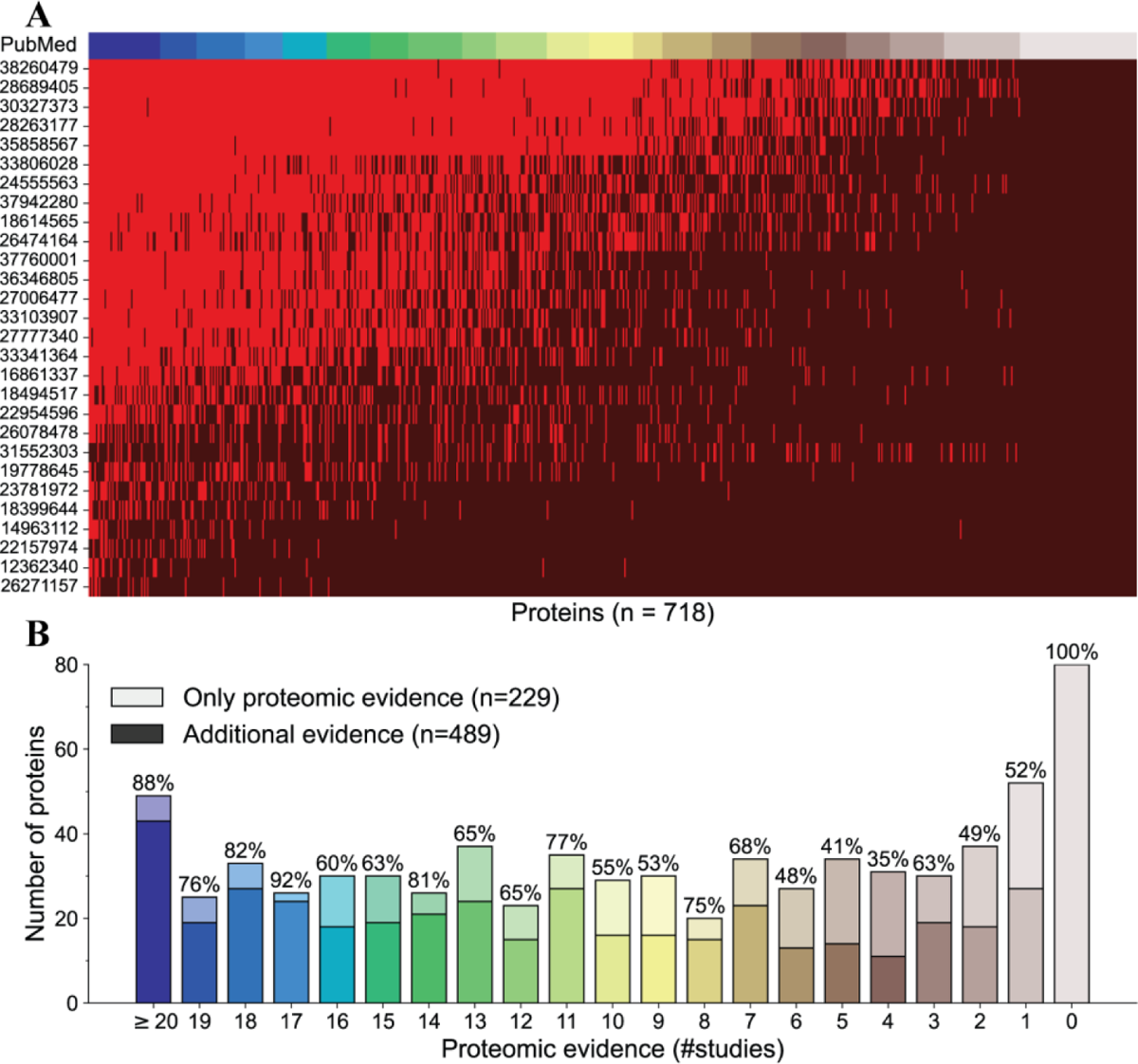
Validation of the RBC-GEM proteome through multi-omic and literature evidence. Proteomic data was collected across 28 datasets to facilitate the development of RBC-GEM. (A) The proteomic evidence for RBC-GEM is visualized as a binary heatmap representing the detection (bright red) or absence (dark red) of proteins across individual studies. (B). Manual curation provided additional evidence for 486 metabolic proteins, with at least 80 proteins not detected in the proteomic evidence.

Proteins that are detected consistently across the 28 proteomic studies were deemed as more likely to be part of the RBC proteome. Furthermore, proteomic technologies have only become more reliable over time[29] by coupling new technologies with emerging methodologies[74] for protein identification[70–73], quantification[27], and purity of mature RBC samples[28]. For this reason, we operated under the assumption that recently published proteomics data are more indicative of the mature erythrocyte proteome compared to extant literature, and prioritized for mapping the collection of recent data sets designed for deep proteome profiling of the RBC cytosol and plasma membrane[26,27,77,78,81,82].

Proteomics data from highly purified reticulocyte populations and erythroid precursors were utilized in an effort to distinguish proteins that could be from a newly matured red blood cell from those that are clearly specific to immature erythroid precursors[28,81]. We also included a high-confidence canonical erythrocyte proteome defined by reconciling proteomic data through supervised machine learning[86]. There is increasing interest in using RBCs within the context of clinical proteomics [91] and transfusion medicine[29]; therefore, we included several studies that explore alterations in the RBC proteome due to various pathological disease states[75,80,83,85,87,88] and refrigerated storage under blood bank conditions[26,84,85], increasing the utility of RBC-GEM for biomarker discovery[43]. Because several proteomic datasets contained outdated and alternate accessions[18,19,70–74,76], the unavoidable consequence of routine database updates and previous database deprecations[92], we consolidated all identifiers into a list and utilized UniProtKB mapping service to update all protein identifiers to current UniProtKB accessions, pruning obsolete and unreviewed proteins from the list. In total, we found that over 4,600 proteins were detected at least once across all collected datasets (Table B in S1 File).

### Metabolomics and literature curation verify enzymatic activity in the low-abundance proteome

While proteins that are detected consistently across studies are deemed to be part of the RBC proteome, it is important to note that the proteome may also include infrequently detected proteins. Differences in MS-instrumentation, fractionation strategies, contamination, and other sources of technical variability can lead to discrepancies in identified proteins. Functional metabolic tracing experiments suggest that there are still proteins in the RBC proteome that have yet to be identified[21,25,26,78]. Additionally, the presence of an enzyme does not necessarily indicate residual activity for that given enzyme. Therefore, both untargeted and tracing metabolomic experiments of human RBCs [4,23,26,44,46–49,93] were utilized to further substantiate the presence of enzymatic activity and validate the existence of related metabolites (substrates/products). Through the use of high-throughput multi-omic data, we were able to gain a comprehensive understanding of the RBC proteome[29,30,94].

There are potential pitfalls that warrant caution when aggregating publicly available proteomic data for the identification of proteins in the RBC proteome. Over 80 proteins were found to have supporting experimental evidence for their presence and activity in RBCs, yet they were not detected across all proteomic studies. For example, at least six different phosphodiesterase proteins have been identified in RBCs with important roles in regulating cyclic nucleotide levels[95], yet none were found in the proteomic data. Proteins involved in the synthesis of blood group antigens also do not appear across proteomic studies[96,97]. Even the most abundant proteins known to exist in the erythrocyte proteome (e.g., hemoglobin and the Band 3 anion exchanger) were not reported across all studies, perhaps because of high abundance protein depletion strategies aimed at unmasking the low-abundance proteome. The absence of evidence for a known RBC protein in a dataset may be caused by technical and methodological reasons, emphasizing the importance of detailed manual curation in the generation of RBC-GEM [4,23,26,39,46–48].

### Properties of the network reconstruction

The properties of the RBC metabolic network are further contextualized through the functional categorization of reactions by their assigned subsystems (Fig. 4). A subsystem is defined as a collection of functional roles that implement a specific biological process or structural complex, and the set of functional roles that tie protein-encoding genes to different subsystems are known as subsystem connections[99]. RBC-GEM contains over 70 subsystems that can be classified into seven distinct metabolic categories, with an additional category representing miscellaneous processes with varying physiological importance (Fig. 4A). Because transport reactions in RBC-GEM are exclusive to one subsystem, we used the Transport Classification System[98] to classify membrane transporters (Fig. 4B). Most reactions found within the RBC-GEM have known gene associations, with a much smaller subset of reactions with known spontaneity (Fig. 4C). Most genes in RBC-GEM had one subsystem connection (Fig. 4E), as is expected of protein-encoding genes[99]. However, genes found in the “Reactive Species” group were distributed across categories with Superoxide dismutase (*SOD1*), Methanethiol oxidase (*SELENBP1*), and sulfiredoxin (*SRXN1*) as notable exceptions. Furthermore, the representative metabolites are found across all aspects of cellular metabolism, hence the “Reactive Species” category containing the largest number of shared metabolites (Fig. 4D) and genes (Fig. 4E). These observations emphasize the need for both reactive species detoxification mechanisms across metabolic subsystems as well as the specific mechanisms for oxidative stress defense and repair[56,100,101].

**Fig 4.**
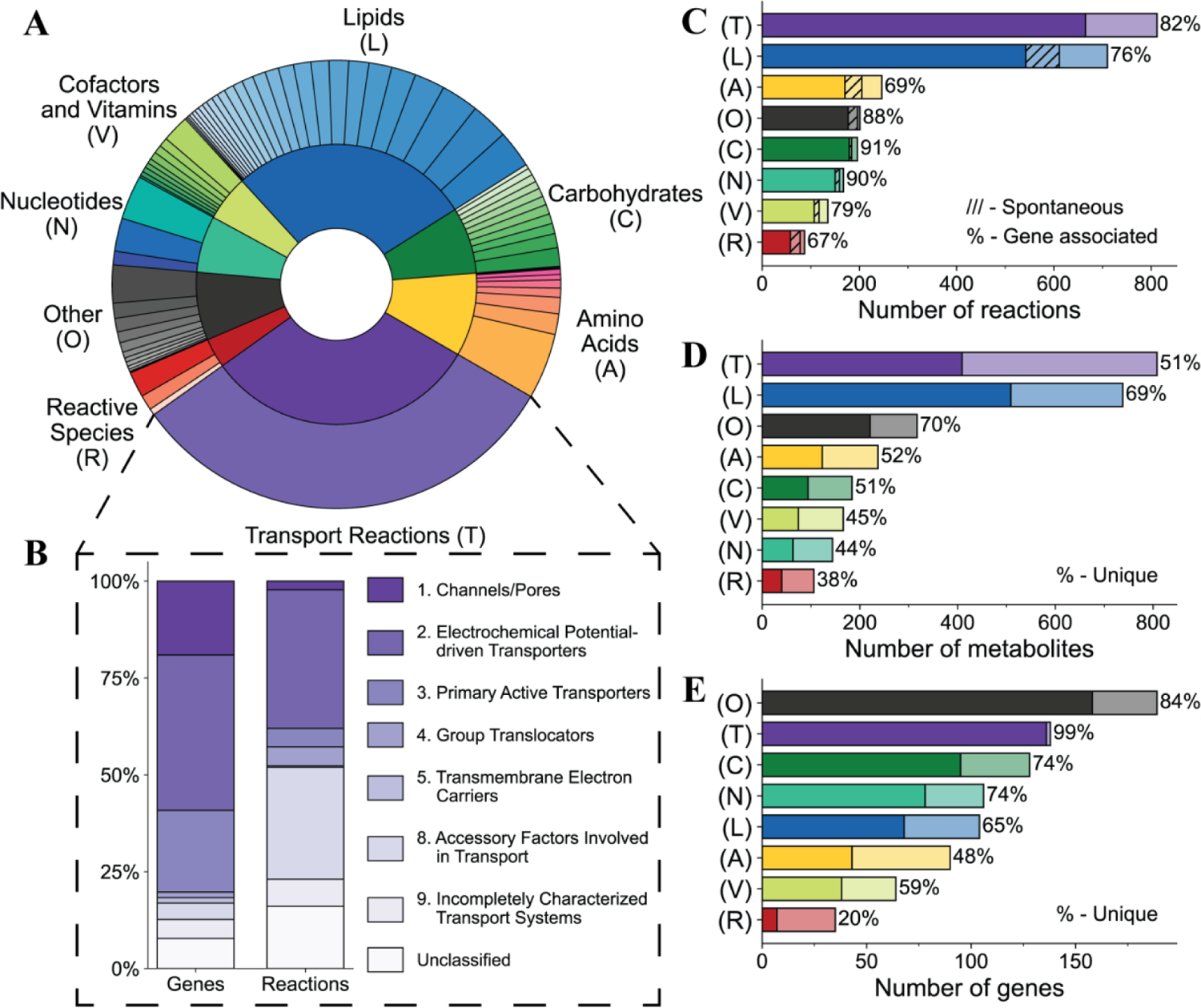
Assessment of RBC-GEM 1.1.0 properties through functional categorization of metabolic subsystems. The various metabolic subsystems represented in RBC-GEM were grouped into eight distinct categories. (A) The relative distribution of 76 metabolic subsystems across the generalized categories. Slice sizes in the inner and outer rings correspond to the number of reactions per category or per subsystem, respectively. (B) Transport reactions across the plasma membrane are further classified according to the Transport Classification System[98]. The “Unclassified” label is assigned to reactions associated with unknown or unclassified transport proteins as well as reactions representing passive diffusion. (C) The number of reactions per category, with darker colored regions representing reactions with gene associations and hashed lighter regions representing known spontaneous reactions without gene associations. (D) The number of metabolites per category, with darker colored regions representing the percentage of metabolites unique to each category. (E) The number of genes per category, with darker colored regions representing the percentage of genes unique to each category. Metabolites and genes are defined as unique to a category if they are exclusively associated with reactions of the same category. The detailed lists of reactions, metabolites, genes, subsystem categories, and transporter classifications are provided in the SI.

### Hemoglobin allostery and pH modulate canonical metabolism

RBCs are well known for their near exclusive reliance on glycolysis for the generation of ATP, the pentose phosphate pathway for NADPH regeneration, and the purine salvage pathway for maintaining adequate nucleotide concentrations. Many of these rate-limiting enzymes are pH sensitive, including hexokinase (*HK1*), phosphofructokinase isozymes (*PFKL*, *PFKM*, *PFKP*)[102], and *G6PD*. The product of glycolysis in RBCs is lactate [31], which exists in equilibrium with lactic acid at physiological pH. The increased lactate contributes to the Bohr effect via intracellular acidification, further promoting the oxygen off-loading from hemoglobin. Furthermore, *CYB5R3* competes with lactate dehydrogenase (LDH) for NADH, a phenomenon contributing to the accumulation of lactate observed in prolonged storage of RBCs[46,103].

The activity of the Rapoport-Luebering (RL) shunt is also pH sensitive and known to dictate the concentration of 2,3-BPG in RBCs. At acidic pH, bisphosphoglycerate mutase (*BPGM*) phosphatase activity is dominant over synthase activity; conversely, at alkaline pH, the synthase activity is preferred. [104]. RBCs also have multiple inositol polyphosphate phosphatase (*MIPP1*) with 2-phosphatase activity for 2,3-BPG [105] and sensitivity to physiologic alkalosis, thus expanding the regulatory capacity of the RL shunt. The changes in 2,3-BPG modulate hemoglobin allostery. The stabilization of deoxyhemoglobin in turn modulates the subcellular compartmentalization of glycolytic enzymes (phosphofructokinase – PFK; aldolase – *ALDOA*; *GAPDH*), which are bound to and inhibited by the N-terminus cytosolic domain of band 3 at high oxygen saturation. At low oxygen saturation, the bound glycolytic enzymes are outcompeted by deoxyhemoglobin and released. The release of glycolytic enzymes to the cytosol corresponds to an increase in the activity of these rate-limiting enzymes the glycolytic pathway[102,106,107], consequently forming an intricate feedback loop in which oxygen transport and delivery is finely regulated through RBC metabolic demands [108].

### Pathological metabolic states elucidate metabolic pathways

RBCs have several proteins, many exclusive to carbohydrate metabolism (Fig. 4D), with metabolic functions for maintaining intracellular energy levels. Though RBCs are capable of transporting many carbohydrates across their membrane[109,110], they best utilize glucose to fuel energy metabolism with approximately 90% of glucose directed to glycolysis and the remaining 10% directed to the pentose phosphate pathway under normal, non-stress conditions[31,111]. Previous metabolomic experiments have revealed the ability of RBCs to metabolize citrate and other tricarboxylic acids[45] in addition to carbohydrates such as fructose, mannose[112], and galactose[113]. Though not immediately visible in healthy individuals, RBCs from individuals with different forms of glycogen storage disease have revealed residual activity of synthesis and degradation enzymes[114]. Proteomic evidence also elucidated multiple metabolite repair enzymes in RBCs that have been shown to address side activities of glycolytic enzymes[115]. Use of this evidence in RBC-GEM is illustrated by the formation of methylglyoxal through oxidation of dihydroxyacetone phosphate (DHAP) [116,117], subsequent detoxification through the glyoxalase enzymes lactoylglutathione lyase (*GLO1*) and hydroxyacylglutathione hydrolase (*HAGH*), respectively known as Glyoxalase I and II[118], and deglycase activity of Parkinson disease protein 7 (*PARK7*)[119].

RBC nucleotide metabolism has been shown to be more complex than previously thought, elucidated through pathological RBC states which are characterized by alterations in nucleotide patterns[120,121]. Nucleotide metabolism was composed of a few metabolic subsystems (Fig. 4A), with nucleotide species found across the RBC metabolite network (Fig. 4E) and regulated by specific enzymes to maintain the nucleotide pools (Fig. 4D). RBCs have enzymes to salvage extracellular orotic acid to form UMP[122], phosphorylation and dephosphorylation of deoxynucleotides[123,124], salvage the nucleobase, and ribose moieties of deoxynucleotides[125]. Erythrocytes have enzymes for uptake of cyclic nucleotides[126,127], signaling[95], degradation[128], and export[126]. Though early studies were unable to detect adenylosuccinate synthetase activity in RBCs[129], deep proteomic and metabolic tracing studies have demonstrated its presence and a residual activity[130]. Furthermore, erythrocytes have the capacity to form unusual nucleotides, including the oncometabolite 4-pyridone-3-carboxamide-1-beta-D-ribonucleotides[131,132] and the Lesch Nyhan (LN) biomarker 5-Aminoimidazole-4-carboxamide ribonucleotides (AICAR)[88,131,132]. LN syndrome is an X-linked recessive inborn error caused by a pathogenic mutation in Hypoxanthine-Guanine Phosphoribosyltransferase (*HPRT1*). *HPRT1* is responsible for salvaging hypoxanthine, a catabolic product of ATP breakdown and deamination [88] by AMP deaminase 3 (*AMPD3*); thus, LN is characterized by urate accumulation due to oxidation of excess hypoxanthine with concomitant generation of reactive oxygen species (ROS). Furthermore, oxidative stress is known to promote *AMPD3* activity while 2,3-BPG inhibits it, thus the modulation of purine metabolism is intertwined with the RBC response to hypoxia via deregulation of *AMPD3* [130,133].

Erythrocytes are constantly exposed to ROS from external and internal sources of oxidants[134], and hemoglobin itself has been shown to participate in a variety of reactions outside of the standard roles it has in gas transport[135–139]. Therefore, we formed the “Reactive Species’’ category with associated subsystems to represent hemoglobin-catalyzed reactions and auto-oxidation events that were likely to occur. Hemoglobins have displayed varying roles in systemic nitric oxide, sulfide, and redox metabolism[56,140]. Furthermore, the degradation of hemoglobin is non-enzymatic, generating reactive species[141] before glutathione facilitates degradation of hemin, as evident in hemolytic disorders like beta thalassemia[142,143]. Because hemoglobin is the most abundant protein in RBCs, we assumed that reactions shown to be catalyzed by hemoglobin *in vitro* were also possible *in vivo* and in blood storage conditions[144–149]. In addition to interactions with hemoglobin, we also included various reactive species detoxification reactions facilitated by intertwined antioxidant networks for “redoxins”, peroxiredoxins[150], thioredoxins[151], and glutaredoxins[152], in which they ultimately derive reducing power from NADPH[56]. The inclusion of several reactions involving the formation and detoxification of reactive oxygen, nitrogen, and sulfur species, especially in the context of their interactions with hemoglobin and “redoxins’’, led to the formation of the “Reactive Species’’ category[140].

### Lipid metabolism in RBCs is complex

Most spontaneous reactions in the lipid metabolism (Fig. 4C) were due to the inclusion of lipid peroxidation reactions, in which hydroxyl species generated by Fenton and Haber-Weiss reactions initiate lipid peroxidation through hydrogen abstraction of representative polyunsaturated lipid species linoleate, arachidonate, eicosapentaenoate, and docosahexaenoate (C18:2, C20:4, C20:5, C22:6), followed by formation of the lipid peroxyl radical and eventual termination by Vitamin E. [56,153,154]. The generation of reactive species through oxidation of catechol estrogens and redox cycling due to *CYB5R3* was also included as estrogen may have profound roles in sex-based differences in blood storage [155–158]. Other updates to lipid metabolism in RBC-GEM reflect the erythrocyte incapacity for *de novo* synthesis of long chain fatty acids due to an inability to maintain a sufficient pool of Malonyl-CoA [159], instead relying on the Lands cycle for phospholipid remodeling and repair[160,161].

Erythrocytes have demonstrated the ability to successively methylate phosphatidylethanolamine (PE) to phosphatidylcholine (PC) via an N-methyltransferase[162], to incorporate glucose, phosphate, glycerol, serine, and choline into phospholipids, and exhibited low phosphatidylserine decarboxylase activity[163]. PC may serve as an unappreciated pool of methyl group donors to fuel protein-L-isoaspartate (D-aspartate) O-methyltransferase (*PCMT1*) and other methyltransferases for deamidation/dehydration-induced isoaspartyl-damage repair, as elucidated through metabolomics of stored RBCs [4,47,53]. Additionally, the abnormal phospholipid phosphatidylethanol was found almost exclusively in RBCs and may serve as a possible biomarker for alcohol consumption[164,165].

Sphingosine-1-phosphate (S1P), the bioactive signaling lipid that modulates the hypoxic response of RBCs[93,166], may experience limited degradation through low-activity of S1P degrading enzymes[167,168], but is primarily exported by the Sphingosine-1-phosphate transporter (*MFSD2B*)[169]. RBCs are capable of efficient sphingosine uptake and phosphorylation to S1P via sphingosine kinase 1 (*SPHK1*) [170], enabling circulating RBCs to effectively serve as reservoirs for plasma S1P. RBCs have also demonstrated alkaline ceramidase, neutral and acid sphingomyelinase, and sphingomyelin synthase activities, indicating that sphingolipid metabolism in the membrane may represent another regulatory point for S1P metabolism[168]. RBC-GEM was updated to include the various metabolic processes for sphingolipid metabolism, thereby representing the processes for erythrocyte metabolic reprogramming due to S1P[171]. Metabolomic studies of RBCs have highlighted consistent sphingolipid phenotypes of neurodegenerative diseases[172] and the role of elevated S1P in promoting sickle cell disease progression[173]. Because RBCs are the largest reservoir of S1P [174,175] and ceramide formation by sphingomyelinase contributes to eryptosis[176], understanding RBC sphingolipid metabolism is essential to elucidate the contribution of RBC-derived S1P in the pathogenesis of various diseases[177]

### RBCs contribute to organismal homeostasis beyond oxygen transport activity

RBC-GEM includes all the known amino acid transporters and the metabolically active enzymes elucidated through proteomics and supported by experimental evidence in the literature. The role of amino acids in erythrocyte metabolism was once thought to be limited to glutathione synthesis[178]; however, proteomic and metabolomic tracing experiments have revealed a diverse set of reactions in erythrocytes utilizing amino acids as substrates to drive metabolic processes. Erythrocytes contain metabolically active enzymes for arginine catabolism in nitric oxide regulation [179–182], transamination for glutamate homeostasis[26,183,184], and methionine salvage[32,185]. The available concentrations of amino acids may modulate the hypoxic response in erythrocytes[44,186]. Consequently, erythrocyte membranes have at least seven different amino acid transport systems that facilitate the rapid exchange of up to 17% of their total amino acid pool with surrounding plasma[187], highlighting their role as circulating reservoirs of amino acids and vitamins for maintaining organismal homeostasis and facilitating cross-talk between RBCs and other cell types[188–190]..

We expanded NAD metabolism with NAD glycohydrolase (*CD38*) activity[54,191,192], dihydronicotinamide riboside (NRH) salvage via adenosine kinase (*ADK*)[193], and oxidation through NRH:quinone oxidoreductase (*NQO2*)[194–196]. We also included *GAPDH* catalyzed and spontaneous generation of hydrated NADH [115], and subsequent repair through enzyme-catalyzed epimerase and ATP-dependent dehydration reactions, once stated to exist in RBCs[197] and later confirmed in approaches to elucidate the RBC proteome[26,27,86]. Erythrocytes have translocator protein 2 (*TSPO2*)[198], involved in cellular import of the heme precursor 5-Aminolevulinic acid[199], cytoplasmic enzymes of the heme biosynthetic pathway (*ALAD*, *HMBS*, *UROS*, *UROD*), and ATP-binding cassette sub-family member 6 (*ABCB6*), involved cellular efflux of porphyrins at the plasma membrane[200,201], thus forming a non-canonical pathway of unknown significance. Pantothenate and CoA metabolism[202,203], folate metabolism with its connections to AICAR metabolism[204,205], thiamine metabolism[206,207], and Vitamin E recycling at the erythrocyte membrane were also included[153,208] in the RBC-GEM network.

RBCs have several other metabolic capabilities that are increasingly being recognized for their physiological and pharmacological relevance. They have been shown to have roles in the endocrine system[209], act as modulators of innate immunity[210,211], and hydrolyze bioactive peptides including Angiotensin II[212–214]. Therefore, we included catecholamines such as epinephrine, norepinephrine, and dopamine along with subsequent oxidation and glutathione conjugation reactions[215]. We also included various post translational modifications and repair reactions, including deglycation by fructose 3-kinase (*FN3K*)[102,216] and related protein (*FN3KRP*)[217], methylation by various methyltransferases including *PCMT1*[185], glycosylation by residual activity of blood group proteins[218,219], signaling through phosphorylation and dephosphorylation[28,220,221], protein degradation through ubiquitin-mediated proteolysis[83,222], and alkylation via transglutaminase (*TGM2*)[223]. The multitude of post-translational modifications (PTMs) and protein repair reactions included in RBC-GEM are therefore indicative of the importance of PTMs for RBC metabolic reprogramming, adaptivity, and timing of senescence in a cell devoid of standard protein synthesis and degradation. PTMs diversify protein functions, therefore understanding post-translational modifications is important to discovering the full functionality of the RBC proteome.

### Classification of membrane transport proteins

Understanding the influence of genetic variation on RBC membrane protein expression is important to understanding pathological metabolic states, exemplified by the formal recognition of several membrane transport proteins as ‘blood group’ proteins due to their immunological and pharmacological significance [224,225]. All Transport reactions in the RBC network were grouped within a single representative subsystem; therefore, we classified transport reactions using the Transport Classification System[98] to assign classification numbers to transport proteins and their associated reactions (Fig. 4B). The “Unclassified” label was assigned to reactions representing passive transport via diffusion in addition to reactions associated with unknown or unclassified transport proteins (Table J in S1 File). Classification of transport proteins and associated reactions revealed that less than 3% of transport reactions in RBC-GEM involve the transport of ions and small molecules through membrane pores and channels. Despite being responsible for a small percentage of reactions, these transport proteins have critical importance for erythrocyte osmotic regulation and gas transport functions; they serve as a nexus for sensing electrical, chemical, and mechanical changes[226–233], dictating the RBC metabolic response accordingly. Proteins in this category include the blood group antigens Piezo-type mechanosensitive ion channel component 1 (*PIEZO1)*, Rhesus complex proteins (*RHAG*, *RHCE*, *RHD*)[229], Aquaporins 1 and 3 (*AQP1*, *AQP3*)[231,234] the urea transporter (*SLC14A1*), as well as Ca^2+^ channel and Ca^2+^-regulated ion channel proteins such as the Gardos channel (*KCNN4*)[227].

The largest class of transporters, both in terms of total number of proteins and in associated transport reaction, was the class of electrochemical potential-driven gradient transporters, with nearly 40% of transport related genes responsible for 35% of the reactions. The majority of reactions represent the different mechanisms of amino acid transport in the RBC; at least seven different amino acid transport systems have been discovered, with unclear metabolic functions in RBCs other than providing the precursors for glutathione synthesis[178]. Further compelling evidence supports the role RBCs play in interorgan amino acid delivery tissues. Thus, the numerous routes for amino acid exchange further highlight the contribution of RBCs toward maintaining metabolic homeostasis outside of their gas transport functions[187,188,190,235,236].

The remainder of reactions reflect carbohydrate uptake[237], monocarboxylate exchange[238], folate exchange[239], facilitated diffusion of nucleotides[120,240–242], ion cotransport and countertransport[232], and the numerous anion exchange reactions facilitated by the Band 3 anion exchanger (*SLC4A1*)[243].

The vast majority of reactions catalyzed by primary active transporters are associated with the ATP-dependent efflux of a broad selection of metabolites, including, but not limited to, phospholipids, porphyrins, xenobiotics, oxidized glutathione, glutathione-conjugated steroids and catecholamines, and lipid peroxidation products. Aside from their broad specificity, the ATP-binding cassette (ABC) transporters form the genetic basis for several known blood groups (*ABCB6*[244], *ABCC1*[245], *ABCC4*[246], *ABCG2*[247]). Conversely, the P-type ATPases in erythrocytes are responsible for the efflux of a small selection of substrates and have significant roles in normal erythrocyte function. The protein complexes within this group catalyze Na^+^/K^+^-ATPase, and Ca^2+^-ATPase activities in the erythrocyte membrane, respectively consuming 40% and 10% of the total ATP produced by erythrocytes to maintain volume homeostasis and the electrochemical gradient across the plasma membrane. Working in concert with the ABC-transporters that “flop” PC, the P4-type ATPases “flip” externalized PE and PS to prolong the overall loss of plasma membrane asymmetry, a key signal for senescence in erythrocytes[62,248]. Erythrocytes also demonstrate V-ATPase activity[249], with recent proteomics suggesting a contributing role in membrane damage through acidification of the extracellular environment of cold-stored RBCs[84].

The smallest classification group of transporters corresponds to electron transport across the plasma membrane. Transmembrane electron transport is facilitated by ascorbate-dependent reductase (*CYBRD1*), responsible for vitamin C homeostasis at the erythrocyte membrane after the maturational loss of the Na^+^-dependent vitamin C transporter[208,250,251]. It should be noted that ferrireductase (*STEAP3*) also belongs to the same class; however, it is primarily an endosomal protein involved in transferrin-mediated iron uptake[252,253]. STEAP3 activity has been measured in mature murine RBCs, whereby it contributes to the reduction of free ferric iron to the reduced ferrous state. In so doing, STEAP3 might promote Fenton chemistry by recycling a rate-limiting substrate for the generation of reactive oxygen species[252].The radical species resulting from excess STEAP3 activity have been linked to elevated lipid peroxidation in mice and humans, a phenomenon associated with RBC vesiculation, splenic sequestration, and extravascular hemolysis. Interestingly, the erythrocyte contains a few proteins with incompletely characterized transport systems, including the non-ABC multidrug exporter, RalA-binding protein 1 (*RALBP1*)[254], the Ca^2+^-dependent phospholipid scramblase (*PLSCR1*)[255], and *TSPO2*, recently shown to mediate 5-aminolevulinic acid uptake[198,199] and VDAC-mediated ATP export[198].

### MIRIAM compliance

There is an increasing consensus among the Systems Biology community that high-quality GEMs are constructed and distributed that adhere to FAIR principles[34,35]. The RBC-GEM was derived from both iAB-RBC-283 (downloaded from the BiGG Models database[40,256], and from Human-GEM (version 1.18.0[38]), downloaded from the MetabolicAtlas[10]. Thus, for the reconstruction to be findable, we utilized unique BiGG identifier standards and supplemented the RBC-GEM with existing MetabolicAtlas annotations where possible. New identifiers were created for reactions and metabolites which had ambiguous BiGG identifiers or were lacking them altogether. Gene identifiers were chosen based on their official symbols as defined by the HUGO Gene Nomenclature Committee (HGNC) database[257], and annotated with corresponding UniProtKB accessions. Unlike iAB-RBC-283, no isoform specificity was assigned to genes so that each gene uniquely represents one current and reviewed entry in the UniProtKB[42]. Through the UniProtKB ID mapping services, we were able to enrich the proteins represented in the reconstruction with compact identifiers for over 60 different databases.

We also enriched RBC-GEM with genetic and pharmacological information by mapping genes a total of 2,244 drugs in DrugBank[258](Table K in S1 File), 4,562 single nucleotide polymorphisms (SNP) in UniProtKB[42] and SNP database[259] (Table M in S1 File), and 594 phenotypes of morbid SNPs from OMIM[260] (Table L in S1 File). Through the MetabolicAtlas annotations, we were also able to enrich metabolites and reactions with the annotations contained in Human-GEM. All annotations adhere to MIRIAM standards[37,261] that can be resolved through Identifiers.org[262]. However, several new additions for RBC-GEM, particularly those classified in the “Reactive species’’ subsystem, are based primarily on RBC-specific literature [56,263] and do not currently exist in the Human-GEM. As our efforts for RBC-GEM 1.1.0 were focused primarily on expanding the proteomic coverage, these new metabolites and reactions were minimally annotated and thus present future areas of opportunity for the refinement of annotations.

### MEMOTE standardization

During the iterative reconstruction process (Fig. 1), the reconstruction was periodically evaluated using the MEMOTE standardized testing suite that carries out quality control tests for metabolic reconstructions [34]. MEMOTE represents a community standard for assessing reconstruction through the application of a series of standardized set of tests and metrics. RBC-GEM demonstrated 100% stoichiometric consistency with all reactions mass-balanced, excluding pseudo-reactions such as boundary exchanges and the few reactions responsible for “pooling” individual lipid species into generic R-groups. Additionally, 99.8% of reactions are charge balanced with the only notable exceptions being the two reactions responsible for the reduction of methemoglobin, by either cytochrome b_5_ or flavin mononucleotide. The inability to balance these reactions is a likely consequence of simplifying the complexity of oxyhemoglobin binding and alpha-beta subunit interactions to a single subunit entity that could be treated as a metabolite within the reconstruction. We utilized the flux balance/variability analysis (FBA/FVA) algorithm implemented in MEMOTE[264] to ensure known and historically important metabolic pathways[54] remained functional. Dead-end metabolites and blocked reactions, especially those with literature evidence indicating functionality in *in vitro* RBCs, were left in the reconstruction as it is often unclear whether proteins are remnants of an imperfect degradation process or exhibit moonlighting functions[86] with potential physiological or pathological significance. The final reconstruction that was generated in this study is thus the largest (in scope and omics and literature coverage), and one that passes all the major QA/QC tests with a MEMOTE score of approximately 80%.

## Conclusion

RBC-GEM 1.1.0 is the most expansive and comprehensive reconstruction of RBC metabolism to date, supported by 28 proteomic datasets (Table B in S1 File) and a bibliome of 1000+ RBC-specific publications (Table C in S1 File). The contents of the reconstruction are made available: genes (Table D in S1 File), metabolites (Table E in S1 File), and reactions (Table F in S1 File), with over 75 years of available research for the human RBC. Biochemical pathways are visualized in the entire RBC metabolic network through a global metabolic network map, generated by the Escher visualization tool. We provide an interactive version of the map, allowing for visual contextualization of different user-defined data. Connectivity analysis demonstrated the RBC metabolic network is not inherently fragmented, and that the achievements made in understanding RBCs through various omics approaches have revealed a richer, more connected metabolic network than previously known. Through the use of high-throughput multi-omic data, a comprehensive understanding of the catalytically active RBC proteome is also achievable[29,30,94]. Furthermore, coalescence of information about pathological metabolic states in RBCs illuminated the physiological significance of ‘moonlighting’ functions of enzymes and previously overlooked pathways.

RBC-GEM is available in a version-controlled GitHub repository (https://github.com/z-haiman/RBC-GEM) where the versions of the reconstruction are hosted as fully annotated SBML files (S3 File) and as as flat table files. Through our implementation of a version-controlled framework, we adhere to principles of data stewardship and ensure that RBC-GEM remains a curated, high-quality knowledge base, consistent with current knowledge and community consensus, as improvements are continually made in a tractable and traceable manner. RBC-GEM represents a critically important step toward the development of a proteomically complete knowledge base for RBC metabolism. This reconstruction paves the way for the development of the next generation of RBC whole cell models.

## Methods

### Preparing iAB-RBC-283 for expansion

RBC-GEM was constructed using the iAB-RBC-283[12] as the initial starting point from which further refinements and expansions could be made using Human-GEM[10] and RBC specific literature. iAB-RBC-283 reconstruction was downloaded from the BiGG Database version 1.6 [265,266] and identifiers were harmonized with the current iteration of the Human-GEM (version 1.18.0 [38]) downloaded from the MetabolicAtlas[267] to assist in providing a common framework to work with both reconstructions.

We addressed the representation of lipids in the RBC metabolic network. Fatty acid chains in iAB-RBC-283 were not represented as generic R-groups as in other human reconstructions, but instead as three individual lipid species. To accommodate for the increased number of lipid species represented in RBC-GEM, we chose to utilize R-groups to represent lipid species in a manner similar to the Human-GEM reconstruction, resulting in the replacement of 125 reactions (Table N in S1 File) and 62 metabolites (Table O in S1 File) in iAB-RBC-283 with 18 pooled reactions and 10 representative species utilizing R-groups. We condensed the carnitine shuttle reactions from three pairs of irreversible reactions with opposite directions into three individual reversible reactions representing the net reaction, justified due to the known reversibility of the carnitine palmitoyltransferase enzyme in RBCs[58]. We also removed 11 intracellular sink and demand reactions to prevent their interference in the identification of blocked reactions and potential areas of reconstruction expansion. iAB-RBC-283 contained 346 protein products, including alternate splice variants, represented by 283 distinct genes. We simplified the gene-protein-reaction associations through the removal of splice variants and associated each gene with both its official gene symbol [257] and UniProtKB identifier, resulting in 283 unique gene entries. The resulting reconstruction, dubbed RBC-GEM 0.3.0, was hereafter treated as the main draft reconstruction for subsequent refinement and expansion.

### Collection of proteomic data

Published proteomic data for erythrocytes was aggregated from 28 datasets[18–20,26–28,70–90] spanning 20 years of RBC proteomic research to facilitate the expansion of the RBC network. iAB-RBC-283 was derived through the integration of three proteomic datasets[18–20] with the global human metabolic reconstruction at the time, Recon1[13]; however, protein databases have since undergone significant changes, including the discontinuation of the International Protein Index (IPI) database in favor of UniProtKB.[92]. Studies utilizing either IPI[18,19,71,72] or GenInfo (GI) identifiers[70,73,74,76] were therefore mapped to UniProtKB identifiers, and the consolidated list of UniProtKB identifiers was run through the UniProtKB ID mapping service[42], ensuring they reflected updated protein entries. Proteins that could not be successfully mapped to the UniProtKB database were confirmed to be obsolete. In total, over 4000 proteins were detected at least once across 20+ different RBC specific proteomic datasets (Table B in S1 File).

### Reassessment of the existing reconstruction

Using the COBRApy python software package (v0.29.0[268,269]), the complete list of 4000+ proteins was first mapped onto the initial RBC-GEM. Genes and biochemicals without proteomic evidence were then subjected to additional scrutiny through targeted literature searches to justify their presence or removal from the reconstruction. Consequently, genes and associated reactions responsible for initial steps of the Kennedy pathway for *de novo* lipid synthesis were removed, as studies demonstrate erythrocytes lack the phosphotransferase enzymes[163], further evident by the accumulation of cytidine phosphodiester compounds in patients with pyrimidine nucleotidase deficiency[270]. Similarly, we also removed the genes and reactions associated with the final steps of heme synthesis as they are known to be localized to the mitochondria; however, steps leading up mitochondrial transport were kept as recent studies demonstrate erythrocytes contain *TSPO2*[198], involved in cellular import of the heme precursor 5-Aminolevulinic acid[199], and *ABCB6*, involved cellular efflux of porphyrins at the plasma membrane[200,201], thus forming a non-canonical salvage pathway of unknown significance. Enzymatic heme degradation was also removed from the reconstruction as evidence demonstrates nonenzymatic heme degradation due to reactive species as the primary route for heme degradation[141]. However, the abundant *BLVRB* was kept for its physiologically important role as the NADPH flavin reductase for methemoglobin reduction in RBCs [271,272]. RBCs lack the capacity for glucuronidation[273] and do not exhibit significant glycerol kinase activity[274]. Erythrocytes have also been shown incapable of significant alpha-glutamyl dipeptide transport[275–278], nor do they have 5-oxoprolinase activity[47]. We therefore removed the reactions and associated genes that enabled these capabilities within the RBC-GEM reconstruction.

The curation process combined with the additional proteomic evidence also led to the replacement of 18 biochemical and transport reactions in favor of analogous reactions with better evidentiary support. Except for mitochondrial enzymes such as coproporphyrinogen-III oxidase (*CPOX*), protoporphyrinogen oxidase (*PPOX*), and ferrochelatase (*FECH*), the vast majority of removed proteins were due to an inability to find direct supporting evidence for their inclusion while evidence was found for other enzymes catalyzing the same activity. We thus removed these genes from the reconstruction; however, we note that as they are removed for a current lack of evidence, additional evidence may call for their reinclusion in a future iteration of RBC-GEM. In summary, 36 distinct biochemical reactions (Table N in S1 File), eight metabolites (Table O in S1 File), and 113 genes (Table P in S1 File) were removed from the RBC-GEM reconstruction.

### Expansion of the RBC-GEM

We mapped the complete list of 4000+ proteins to the Human-GEM reconstruction to identify candidate reactions to add to the RBC-GEM.[279,280]. After filtering out the reactions occurring within organelles, we explored the remaining candidate reactions in a subsystem-dependent manner. For each subsystem, we started with reactions associated with proteins that appear in multiple proteomic datasets, and searched the literature for supporting evidence. Often, we were able to find supporting evidence in the form of human RBC-specific literature; however, we relied on biochemical databases (e.g., KEGG[41], UniProtKB[42] RHEA[281]), pharmacological databases[282], and other literary sources pertaining to other human cell types or erythrocytes in similar species. We also relied on literature on RBC pathological states to verify typically inactive or non-canonical pathways in the RBC; for example, pyrimidine 5’-nucleotidase deficiency[124] provided insight into deoxynucleotide detoxification. As a cell without protein synthesis capabilities, the presence and relevance of enzymatic activity in RBCs is often subject for debate; therefore, we included all evidence utilized in the curation process for addition of genes (Table D in S1 File) and reactions (Table F in S1 File).

Erythrocytes are constantly exposed to ROS from external and internal sources of oxidants[56]; therefore we made the assumption that spontaneous oxidation events due to reactive species are more likely to occur than other types of reactions. Additionally, hemoglobin itself has been shown to participate in a variety of reactions outside of the standard role it has in gas transport[135–139]. Furthermore, the degradation of hemoglobin is non-enzymatic, generating reactive species[141] before glutathione facilitates degradation of hemin, as evident in hemolytic disorders like beta thalassemia[142,143]. Hemoglobin is the most abundant protein in red cells, we therefore assumed that reactions shown to be catalyzed by hemoglobin *in vitro* were also possible *in vivo* [144–148]. The inclusion of several reactions involving the formation and detoxification of reactive oxygen, nitrogen, and sulfur species, especially in the context of their interactions with hemoglobin, led to the formation of the “Reactive Species’’ category.[140].

### Curation of subsystem categories

Assignment of reactions to metabolic subsystems can provide meaningful biological context within a larger network analysis, aiding in their analysis and visualization [51]. We therefore assign subsystems following conventions used in the Human-GEM[10]. The subsystems were then compared to KEGG database[41] in order to broadly group subsystems into the general categories “Amino acid metabolism”, “Carbohydrate metabolism”, “Lipid metabolism”, “Metabolism of cofactors and vitamins”, and “Nucleotide metabolism”. Subsystems pertaining to both hemoglobin and spontaneously formed reactive species were categorized as “Reactive species”. All other subsystems, such as those pertaining to post-translational modifications of proteins or peptide metabolism, were categorized as “Other” for clarity in visualization (Table G in S1 File).

### Visualization of the global RBC metabolic network

In tandem with the development of RBC-GEM, we developed a network map of the full erythrocyte metabolic network using the Escher Network visualization tool[50]. In the construction of the map, we grouped reactions within the context of their subsystems and subsequently color-coded reactions based on their generalized category. We focused our initial efforts on the development of a global metabolic map of erythrocyte metabolism so that users of RBC-GEM could contextualize relevant information within the full network context before tailoring the reconstruction for specific applications. The map is provided as an Escher JSON file, enabling users to apply it for their own purposes, including the derivation of new pathway visualizations via Escher without having to start from a blank canvas. The metabolic map is also provided in standard SBGN and SBML layouts, generated by EscherConverter tool (version 1.2), for accessibility and interoperability with other network visualization tools. All files are found in the supplement (S2 File).

### Functional testing of metabolic capabilities

Throughout each iteration of the reconstruction processes, the COBRApy implementation of flux variability analysis (FVA)[264,283] was utilized to calculate the minimum and maximum allowable flux through each reaction. Reactions with non-zero flux values may be activated under specific conditions, indicating that they may have relevant physiological functions. For reactions producing a zero flux, literature was consulted to determine whether the associated pathways should be functional in the erythrocyte. As an example, erythrocytes have demonstrated capability to synthesize and elongate lipids so long as Malonyl-CoA is present in the environment; it is the inability for erythrocytes to produce Malonyl-CoA at a non-negligible rate that prevents activation of the pathway. Consequently, all reactions that stem from Malonyl-CoA produce a zero flux in FVA simulations. Another example can be seen in the treatment of the acyl-carnitines throughout the reconstruction. Acyl carnitines serve as markers of membrane integrity in erythrocytes[83,161,284,285]; however, the maintenance of the pool involves the reversible transfer of the acyl group between carnitine and coenzyme A. Consequently, these reactions require the addition of pseudoreactions to produce a non-zero flux in FVA simulations. Therefore, we opted to keep these reactions and metabolites in reconstruction regardless of the flux value produced by FVA; empowering users of RBC-GEM to make their own decisions regarding relevance and inclusion in their applications.

### Calculation of metabolite and gene connectivity

The stoichiometric matrix of RBC-GEM was used to compute the metabolite connectivity for each metabolite, defined as the total number of reactions in which the metabolite participates in. After removing all pseudoreactions representing system boundaries and the pooling of lipid species, the connectivity of each metabolite was calculated through summation of the representative row in the stoichiometric matrix. Metabolite connectivities were determined with and without partitioning species by compartments. Resulting connectivities were rank ordered, forming a discrete distribution for subsequent comparison to the calculated connectivity distribution for iAB-RBC-283.The gene connectivity was defined as the total number of reactions associated with the gene. The number of reactions per each gene was summed and the results were subsequently rank ordered.

### MIRIAM compliance and MEMOTE standardized testing

Recon3D[9] downloaded from the BiGG Database version 1.6[265,266] was used throughout the reconstruction process to map between BiGG and MetabolicAtlas identifier namespaces. Newly added reactions and metabolites that were unique to RBC-GEM were given new identifiers following BiGG ID standards. Reactions and metabolites previously assigned ambiguous or non-descriptive BiGG identifiers were also assigned new identifiers following BIGG standards following the retirement of their original ID within the RBC-GEM. New reactions were mass balanced using the Chemicalize application [286] to compute the molecular formula and charge of associated metabolites at pH 7.25. The species representative of protein residues and hemoglobin subunits in RBC-GEM included generic R-groups in their formula to represent the remainder of the macromolecule after mass and charge-balancing associated reactions.

Through the MetabolicAtlas, we enriched metabolites and reactions with the connections to external resources originally contained within the Human-GEM. Rather than adhere to the use of Ensembl identifiers as seen in Human-GEM, we opted for gene identifiers based on the HUGO Gene Nomenclature Committee (HGNC) database as we felt they were human readable, short and memorable and SBML compliant in most circumstances. Therefore, each gene/protein contained in RBC-GEM was uniquely identified by its gene symbol[257] and set to represent one reviewed and current entry in the UniProtKB database[42]. We utilized the UniProtKB ID mapping service to extract external identifiers connecting gene information across multiple resources and annotated RBC-GEM accordingly. Compact identifiers for over 60 different databases could be extracted and mapped into the RBC-GEM, enriching protein in the reconstruction with compact identifiers from over 60 different databases. We also mapped the proteins in RBC-GEM to other databases (Table A in S1 File) for pharmaceutical and genetic information, including drugs found in DrugBank [258] (Table K in S1 File), phenotypes in OMIM[260](Table L in S1 File), and SNPs found in the UniProtKB[42] and SNP databases[259](Table M in S1 File). All annotations adhere to the guidelines for minimum information requested in the annotation of biochemical models (MIRIAM)[37,261] with compact identifiers that can be resolved through Identifiers.org[262].

MEMOTE (version 0.17.0) was used to highlight changes made and identify knowledge gaps throughout the reconstruction processes. We conFigd the version-controlled GitHub repository to run the MEMOTE standardized test suite with the use of GitHub Actions at each main deployment. Thus through the combination of MEMOTE and GitHub, we ensured RBC-GEM is distributed within a framework for continuous integration of updates and delivery of the latest, quality-controlled version of RBC-GEM.

## Acknowledgements

The authors would like to acknowledge Marc Abrams for providing feedback on the manuscript. Created with BioRender.com: Fig 1, Fig 5A.

## Supporting information

**S1 File.** Supplementary tables. A table of contents containing titles and legends is included on the first sheet.

**S2 File.** Different file formats for the RBC-GEM metabolic network map. The map is represented as a browsable HTML file, a JSON file for Escher, a PDF file, and both SBGN and SBML formats generated using EscherConverter 1.2.

**S3 File.** The fully annotated RBC-GEM reconstruction in Systems Biology Markup Language. The fully annotated RBC-GEM reconstruction is provided in Systems Biology Markup Language.

